# SMORES: A Simple Microfluidic Operating Room for the Examination and Surgery of *Stentor coeruleus*

**DOI:** 10.1101/2024.02.05.578956

**Authors:** Kevin S. Zhang, Ramon Rodriguez, Sindy K.Y. Tang

## Abstract

Ciliates are powerful unicellular model organisms that have been used to elucidate fundamental biological processes. However, the high motility of ciliates presents a major challenge in studies using live-cell microscopy and microsurgery. While various immobilization methods have been developed, they are physiologically disruptive to the cell and incompatible with microscopy and/or microsurgery. Here, we describe a Simple Microfluidic Operating Room for the Examination and Surgery of *Stentor coeruleus* (SMORES). SMORES uses Quake valve-based microfluidics to trap, compress, and perform surgery on *Stentor* as our model ciliate. Compared with previous methods, immobilization by physical compression in SMORES is more effective and uniform. The mean velocity of compressed cells is 24 times less than that of uncompressed cells. The compression is minimally disruptive to the cell and is easily applied or removed using a 3D-printed pressure rig. We demonstrate cell immobilization for up to 2 hours without sacrificing cell viability. SMORES is compatible with confocal microscopy and is capable of media exchange for pharmacokinetic studies. Finally, the modular design of SMORES allows laser ablation or mechanical dissection of a cell into many cell fragments at once. These capabilities are expected to enable biological studies previously impossible in ciliates and other motile species.

## 1. INTRODUCTION

Ciliates, including *Tetrahymena* and *Paramecium*, are powerful unicellular model organisms that have been used to uncover fundamental insights in diverse biological processes from DNA replication to the microtubule cytoskeleton^1–3^. Recently, the ciliate *Stentor coeruleus* has received renewed interest as a model for studying wound healing, regeneration, habituation and learning in a unicellular system^4–7^. Nevertheless, beyond a few select species, ciliate biology has remained largely unexplored, in part due to technical challenges in studying these organisms. Compared with mammalian cells and other popular model systems, the tools available for studying ciliates are much more limited. While fluorescent staining or electron microscopy of fixed ciliates have revealed details of cellular structure and organization, they cannot be used to study dynamic processes in live cells. Manual microsurgery of ciliates, typically performed by hand with a glass microneedle, has long been used to study phenomena such as wound healing and regeneration in *Stentor*^4,8–10^. However, manual surgery is slow, tedious, and lacks reproducibility. To realize the rich potential in ciliate models, there is a crucial need for advanced live-cell microscopy and microsurgery techniques.

Just like any biological sample (i.e., the cell or organism to be studied), a ciliate sample must be first immobilized to achieve high temporal and spatial resolution in live-cell microscopy and microsurgery. Immobilization is especially challenging for ciliates because of their high motility, powered by cilia distributed throughout the cell surface^11^. Observations of ciliates using light microscopy can also be challenging because some species exhibit avoidance behavior upon light stimulation in the visible spectrum^11^. Multiple techniques have been developed to immobilize ciliates and other motile organisms such as C. elegans and various protozoa^12,13^, but these techniques carry major biological or technical limitations. Chemical and drug treatments have been used to paralyze the sample, but they may be cytotoxic and could disrupt physiological processes^4,14,15^. Tuning conditions of the external environment such as CO_2_ levels or temperature have also been used, but they may be similarly disruptive physiologically^4,16–18^. Physical inhibition such as adhering the sample to a substrate, embedding the sample in a gel, and increasing the media viscosity are typically minimally invasive, but sample retrieval can be difficult^19–21^. Another physical inhibition technique involves feeding the sample magnetic particles and using a permanent magnet to hold the ingested particles and thus the sample in place^22^. While the magnet can be removed quickly to release the immobilization, the ingested magnetic particles can obstruct imaging of the internal details of the sample.

Physical immobilization by mechanical compression has distinct advantages over the aforementioned techniques. If designed properly, mechanical compression is minimally disruptive, can be applied and removed quickly, and allows for simple sample retrieval. Three major approaches to immobilization by mechanical compression have been developed. A first, relatively simple approach immobilizes samples by compressing them to a fixed thickness between two substrates. The thickness can be set, for example, by microbead spacers or by patterned PDMS structures^23–26^. While this approach is simple to use and can be scaled readily to multiple samples, the degree of compression cannot be adjusted dynamically. Therefore, cells or organisms of heterogeneous sizes will not be immobilized to the same extent. A second, more adaptive approach immobilizes samples using a mechanical microcompressor^27,28^. In the microcompressor design, two rigid substrates are mounted to threaded parts that can be screwed closer to or farther from each other, thereby controlling the degree of compression to within a few micrometers. The microcompressor was compatible with microscopy and has been modified to incorporate features such as media perfusion. However, the microcompressor requires bulky precision-machined parts to control the compression, which limits its ability to scale to multiple samples. Furthermore, the microcompressor is unlikely to be compatible with microsurgery experiments since the relative rotation of the substrates during device operation could introduce undesirable shear stress and possible damage to the sample. A third approach immobilizes samples using microfluidic Quake valves^29–34^. The Quake valve design consists of a thin PDMS membrane sandwiched between a control channel and a flow channel containing the sample.

Pressurizing the control channel deforms the membrane to compress and immobilize the sample in an adjustable manner. The microfluidic basis of such devices enables high scalability and throughput of experiments. However, conventional Quake valve design immobilizes a sample by pushing it into the corner of a rectangular flow channel. In the corner, the membrane deflection is limited by the channel sidewall, and the cross section of the immobilized sample becomes roughly wedge-shaped. Thus, a conventional Quake valve design is not suitable when a high degree of flatness is required in imaging or manipulating the sample.

In this paper, we describe the design, fabrication, and application of a Simple Microfluidic Operating Room for the Examination and Surgery of *Stentor coeruleus*, referred to as SMORES. The core of the SMORES platform is a microfluidic cage trap comprising a modular poly(dimethylsiloxane) (PDMS) enclosure. The ceiling of the enclosure, containing patterned features, is deformed using Quake valve-based microfluidics to immobilize a sample by mechanical compression. The cage trap serves as an “operating room” to trap, immobilize, and perform cell surgery on *Stentor coeruleus* as our model organism. Compared with conventional Quake valve design, our design achieves a highly uniform compression across the cell area. The use of a hand-operated 3D-printed pressure rig allows simple control of sample compression from ∼20 µm to 100 µm. Just like previous compression-based immobilization methods, our approach is minimally disruptive to the cell and is reversible. The operation of the system – loading, immobilization, and retrieval of samples – is simple and fast (typically < 1 minute per step). For longitudinal studies, the cage trap retains the sample inside the trap even when the trap is uncompressed. Additionally, it is straightforward to exchange the media inside SMORES. Finally, it is simple to modify the modular design for desired applications. We demonstrate the use of SMORES for confocal imaging of *Stentor*, laser ablation of the cell surface, and mechanical dissection of the cell into multiple fragments at once.

## 2. RESULTS AND DISCUSSIONS

### Design of the SMORES platform

The SMORES platform was a microfluidic cage trap system consisting of a cage trap and a control channel (Fig. 1 and Supplementary Fig. S1). These components were arranged across two PDMS layers – a control layer seated above a flow layer bonded to a glass coverslip (Fig. 1a). The operation of the platform was similar to that of a conventional Quake valve, where pressurization of the control layer deformed the thin ceiling of the flow layer.

**Figure 1.**
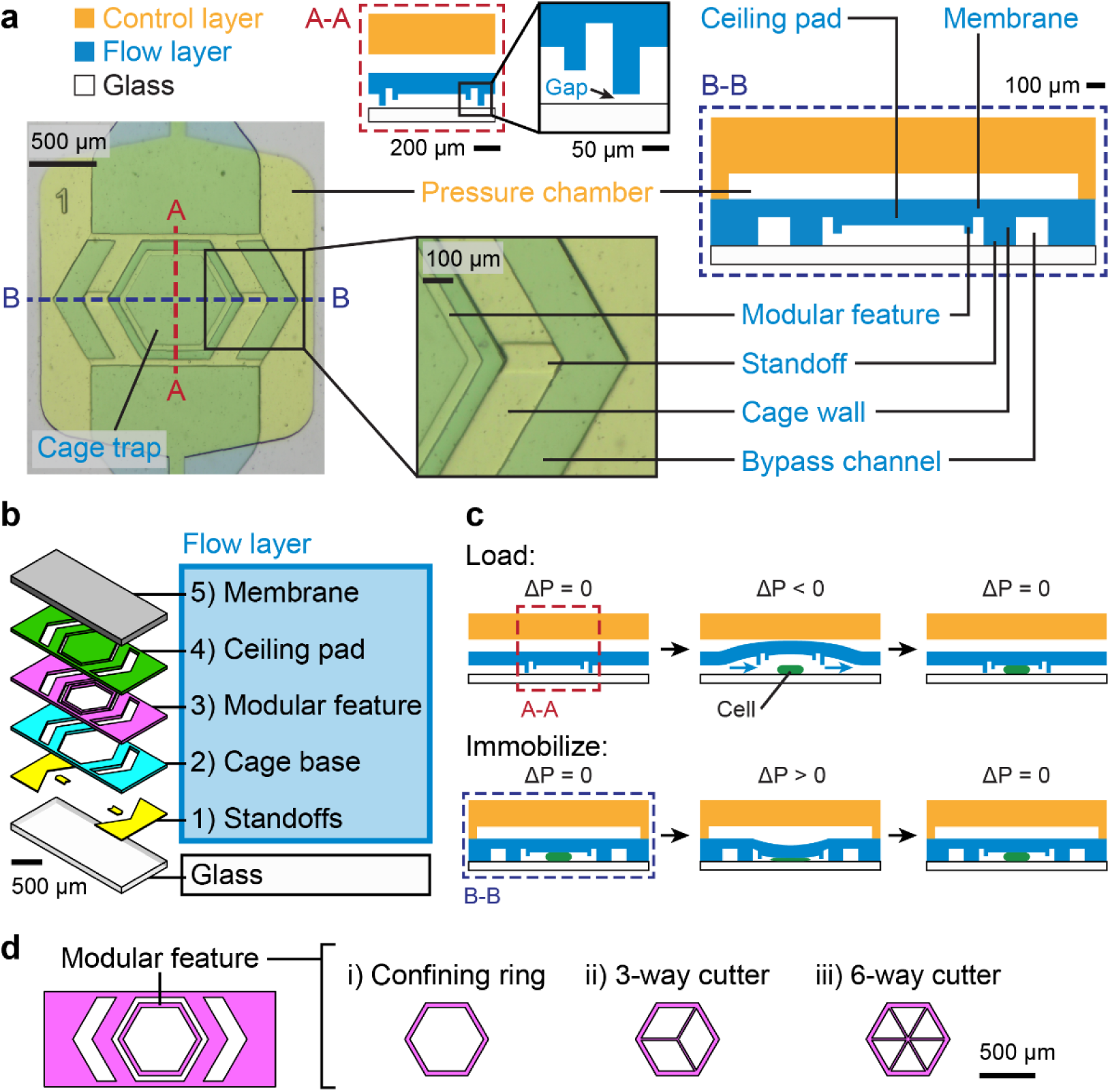
Design of the SMORES platform. (**a**) Left: Top view of a device filled with food coloring indicating the control layer (yellow) and the flow layer (blue). Right: Schematic diagrams of cross sections A-A and B-B corresponding to the dashed lines in the top view. (**b**) Exploded view of the layered components of the flow layer plus the glass coverslip. (**c**) Process flow to load cells (top) and immobilize cells by compression (bottom). ΔP indicates the pressure difference of the control layer relative to the flow layer and is controlled by a 3D-printed pressure rig. Cross sections A-A and B-B are as shown in (**a**). (**d**) Modular feature designs: i) confining ring, ii) 3-way cutter, and iii) 6-way cutter.

The flow layer consisted of a cage trap connected to an inlet and outlet for flowing cells and media. The cage trap was formed by 5 layered components: 1) standoffs, 2) cage base, 3) modular feature, 4) ceiling pad, and 5) membrane (Fig. 1b and Supplementary Fig. S1).

Standoffs and a cage base formed the bottom of the vertical cage wall. The vertical cage wall enclosed the cage trap and was ∼130 μm tall, with a thin gap (∼15 μm) at the bottom due to the standoffs, which were the only features bonded to the glass substrate (Fig. 1a and Supplementary Fig. S1). This gap height (i.e., the standoff height) was chosen to be small enough so that the cells, which were constrained to ∼100 μm in height by the uncompressed cage trap, could not escape through the gap when the trap was uncompressed (100% trap occupancy at 24 hours, n=32/32 cells). On the other hand, the gap height was large enough so that the standoffs prevented the trap from completely sealing when compressed. Since the deflection of the cage trap was constrained near the standoffs, the space about the standoffs remained open to fluid flow even when the cage trap was compressed. This design was critical because if the cage trap was completely sealed against the glass coverslip, fluid would not be able to flow out of the trap (i.e., the trap volume would remain constant). In this case, further compression of the trap would be impossible, and the immobilization of the cell would be poor.

The ceiling of the cage trap was formed by a modular feature that could be modified depending on experimental needs, a ceiling pad that acted as a proof mass to ensure more uniform compression, and a ∼100 μm-thick membrane that separated the flow layer from the control layer (Fig. 1a and Supplementary Fig. S1). To facilitate the loading of a cell into the cage trap by hand and avoid unintentional wounding of the cell, we placed bypass channels on both sides of the cage trap (Fig. 1a). The bypass channels increased the total cross-sectional area of the channel to reduce the flow velocity through the cage trap, but they were too small for cells to enter. The bypass channels also enabled the exchange of media in the trap without accidental loss of the cell.

The control layer consisted of a pressure chamber, a damper, an inlet, and an outlet (Fig. 1a and Supplementary Fig. S1). The pressure chamber was positioned directly above the cage trap. To pressurize the control channel and actuate the cage trap, we built a simple, manually controlled pressure rig consisting of a 3D-printed leadscrew setup holding a syringe filled with cell media (Supplementary Fig. S1). To deform the cage trap towards or away from the glass coverslip, we applied a positive or negative pressure to the control layer by screwing the pressure rig clockwise or counterclockwise, respectively. To absorb any undesirable transient pressure fluctuations, we placed a damper between the pressure rig and the pressure chamber. The damper absorbed pressure changes because it was much larger than the pressure chamber and was the first to deflect when pressure was applied to the control channel. Without the damper, the pressure chamber would be pressurized when the connection tubing was connected or disconnected from the device and could damage the cell inside the cage trap. To provide space for the damper to deflect, we placed a large cavity open to the ambient atmosphere in the flow layer directly underneath the damper.

To load the cells, we lifted the cage trap away from the glass coverslip and then flowed cells into the cage trap from the flow layer inlet by hand (Fig. 1c). We lifted the cage trap sufficiently high (i.e., the gap at the bottom increased to ∼50 µm) so that cells (∼100 µm in height) were not damaged as they flowed under the cage wall. After we flowed a cell into the cage trap, we lowered the cage trap back to a neutral position to trap the cell. To immobilize the cells for imaging or other operations, we applied positive pressure to compress the cell (Fig. 1c). After imaging and/or other operations, we raised the cage trap slowly away from the cell back to a neutral position. The cell could either be left inside the uncompressed cage trap for observation or be retrieved for analysis.

In this work, we tested 3 different modular features (Fig. 1d). For imaging and laser ablation of the cell, we used i) a confining ring that enhanced cell immobilization. For mechanical dissection of the cell, we combined the confining ring with either ii) a 3-way cutter or iii) a 6-way cutter.

### Characterization of cage trap compression

We first characterized the degree of compression achieved by SMORES. We controlled the deflection of the cage trap ceiling by manually dialing the leadscrew of the hand-operated pressure rig. Simultaneously, we measured the ceiling deflection using confocal cross sections of a device filled with fluorescein isothiocyanate (FITC) as a contrast dye (Fig. 2a). The profile of the deflected ceiling was roughly a circular arc. The minimum height *y* of the trap, defined as the distance between the glass and the center of the deflected ceiling (i.e., the arc), decreased from ∼100 to 20 µm approximately linearly with leadscrew rotation (Fig. 2b). The chord (width) of the arc (∼800 µm) was >10x larger than the height (∼0 – 80 µm) of the arc. For this geometry, since the control layer was filled with an incompressible fluid, the linear displacement of the syringe by leadscrew rotation resulted in a linear volume change in the control layer that contributed to the deflection of the ceiling and a linear change in *y*.

**Figure 2.**
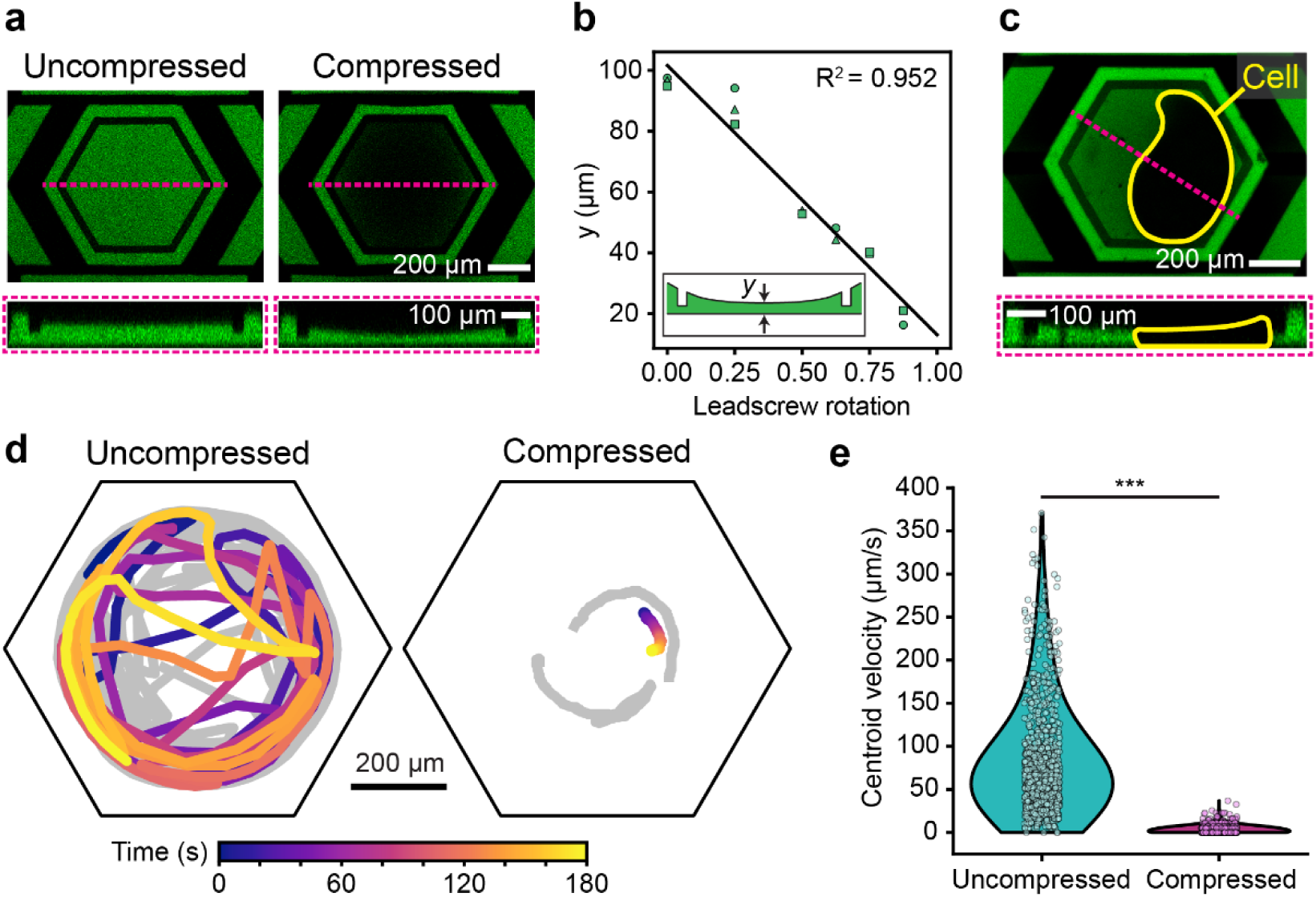
Characterization of cage trap compression. (**a**) Representative confocal images of an uncompressed vs. compressed cage trap showing the deflection of the cage trap ceiling when compressed. (**b**) Minimum cage trap height *y* as a function of leadscrew rotation (linear fit, R^2^=0.952). Inset illustration shows the definition of *y*. Marker shapes correspond to data from independent devices (n=3 devices). (**c**) Representative confocal images of a compressed *Stentor* cell (yellow outline). Cage traps in (**a**) and (**c**) were filled with FITC (green) for contrast and cross sections are indicated by the pink dashed line. (**d**) Motility tracks of cells inside uncompressed vs. compressed cage traps (n=3 cells per condition). One representative track per condition is shown in color and additional tracks are shown in gray. Each track represents 3 minutes of observation. Cage trap walls are indicated by black outlines. (**e**) Centroid velocities of cells inside uncompressed vs. compressed cage traps, shown as violin plots overlaid with individual data points (n=3 cells per condition) (***: P<0.001, two-sample Kolmogorov-Smirnov test).

We then demonstrated that SMORES immobilized *Stentor* cells successfully (Fig. 2c, d, e). In our early prototypes without the confining ring (i.e., a conventional Quake valve design), compressing the trap would typically push the cell against the cage wall where the ceiling deflection was minimal and non-uniform. We observed that a trapped *Stentor* cell could swim towards the cage wall and thus escape from the compression within 1 – 2 minutes. To prevent this behavior, we added a confining ring (Fig. 1d) which followed the perimeter of the ceiling pad and protruded ∼45 µm downwards from the bottom surface of the pad (Fig. 1a). The confining ring retained the cell close to the center of the trap and achieved a compression that was both greater and more uniform than that without the confining ring (Fig. 2c). With the confining ring, the height of the compressed cell varied from ∼30 µm at the center of the trap to ∼60 µm right inside the confining ring (i.e., a 100% variation relative to the height at the center). This variation was due to the circular profile of the ceiling deflection (Fig. 2c). Despite this variation in height, the bulk of the cell was compressed relatively uniformly to ∼30 µm (Fig. 2c). Without the confining ring, the height of the cell would be ∼150 µm at the cage wall (i.e., a 500% variation). The confining ring prevented the cell from escaping from the compression (i.e., into the region between the confining ring and the cage wall) for 45 minutes (n = 4 cells), and up to 2 hours (n = 2 cells). Such extended periods of compression did not compromise cell viability (100% viability 24 hours after release, n=4/4 cells).

To quantify the degree of immobilization, we compared the motility of cells in uncompressed vs. compressed devices. When compressing a cell, we simultaneously monitored the cell to avoid excessive compression and ensure no damage occurred. Damage to the cell appeared in brightfield microscopy as ruptures in the characteristic striped patterns of pigment granules and microtubule ribbons (km fibers) located in the *Stentor* cortex. Motility tracks of the cells showed that uncompressed cells exhibited high motility, while compressed cells were immobile (Fig. 2d). Further, the mean centroid velocity of compressed cells (3.3 µm/second) was almost 24x lower than that of uncompressed cells (78.2 µm/second), and the difference between the two velocity distributions was statistically significant (P<0.001, two-sample Kolmogorov-Smirnov test) (Fig. 2e, Supplementary Fig. S2, and Supplementary Video S1 and S2).

### Confocal imaging of live *Stentor* using SMORES

Because *Stentor* is highly motile, confocal imaging of a live *Stentor* cell is impossible without immobilization. Since SMORES successfully immobilized the cell, we could easily acquire confocal z-stacks of the cell in multiple fluorescent channels. As a proof of concept, we acquired a confocal z-stack of an immobilized *Stentor* cell stained for lipids (DiD and Nile Red) and DNA (Hoechst) (Fig. 3 and Supplementary Video S3). The z-stack took ∼30 seconds during which cell motion was not observable at the resolution of our imaging system. The images revealed the macronuclei, intracellular lipids (e.g., individual vesicles), and cell membrane clearly, and are comparable in quality to images obtained in cells immobilized by the microcompressor^27,28^.

**Figure 3.**
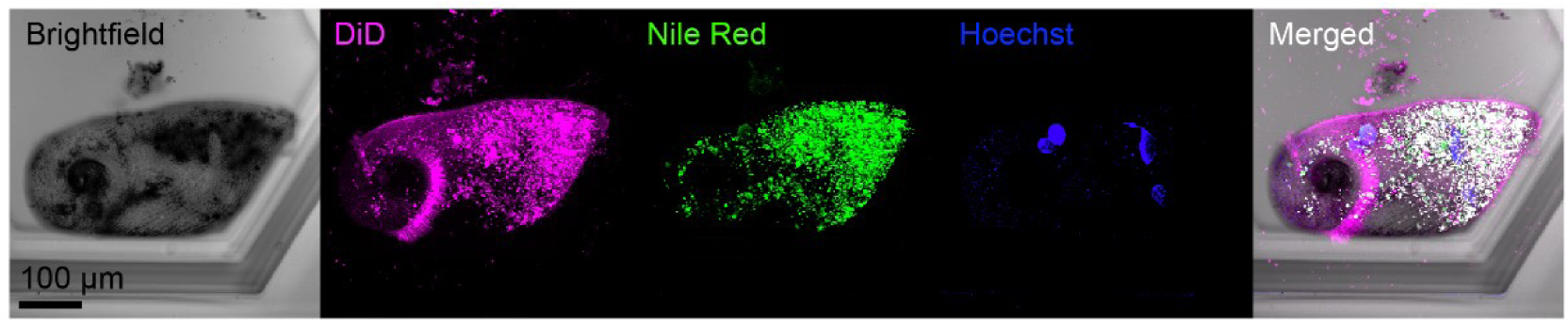
Confocal imaging of live *Stentor* using SMORES. Brightfield, maximum intensity projections, and merged images shown for a representative cell stained for lipids (DiD, magenta; Nile Red, green) and DNA (Hoechst, blue).

### Media Exchange using SMORES

Next, we demonstrate that SMORES allowed simple exchange of media inside the cage trap while retaining the cell inside. We exchanged the media in 3 steps – (1) we compressed the cell inside the cage trap to a thickness of ∼30 µm at the center of the trap, (2) flowed the desired media by hand into the flow layer inlet, and finally (3) released the compression on the cell. To retain the cell inside the cage trap during this process and to avoid damaging the cell, it was crucial to compress the trap before flowing the desired media. For a compressed trap, the cage walls sealed against the glass (except in a small region about the standoffs) and thus isolated the contents of the cage trap from the surrounding media. Therefore, when we introduced the desired media into the flow layer, the media flowed through the bypass channels on either side of the sealed cage trap and did not flow through the trap itself. This operation limited the risk of damaging the cell by shear or accidentally pushing the cell out of the trap. Once the desired media fully surrounded the sealed cage trap, we released the compression of the trap. The desired media outside the trap was allowed to mix with the media inside the trap by diffusion. We demonstrated this process by exchanging deionized (DI) water inside the cage trap with a fluorescent dye solution (5 µM FITC in DI water) (Fig. 4a). This experimental setup allowed us to assess the speed of media exchange within the trap. The mean fluorescence intensity inside the cage trap increased to the reference intensity, as measured by the fluorescence intensity of the dye solution in the channel outside the cage trap, within 35 seconds after we released the compression on the cage trap (Fig. 4b).

**Figure 4.**
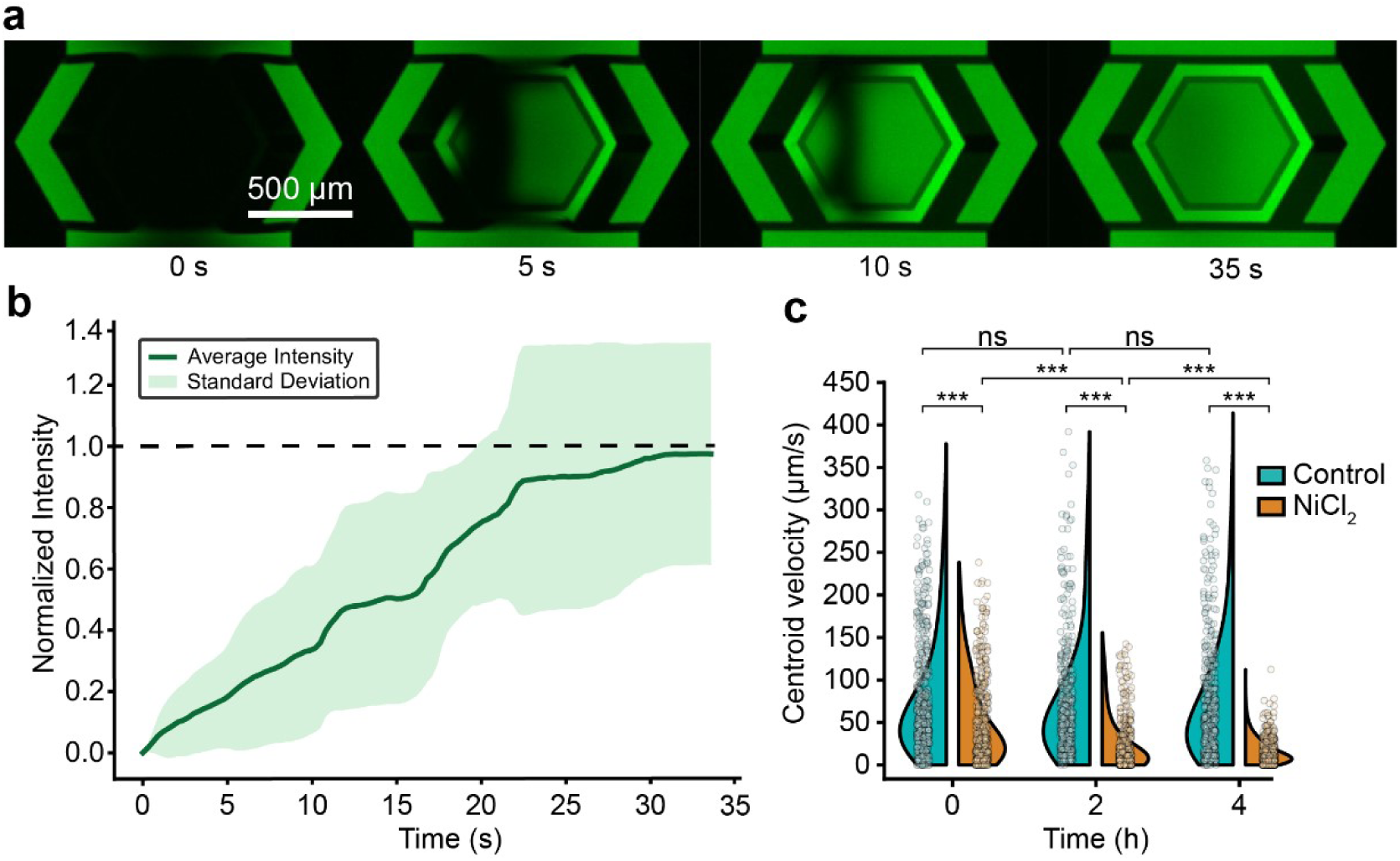
Media exchange using SMORES. (**a**) Time series showing the influx of FITC into the cage trap after release of compression. (**b**) Intensity plot measuring the mean fluorescence intensity inside the cage trap normalized to the reference fluorescence intensity of FITC solution in the channel outside of the cage trap (n=9 devices). Dashed line indicates normalized reference fluorescent intensity of 1. (**c**) Centroid velocities of untreated control cells and NiCl_2_-treated cells inside uncompressed cage traps over time. Data for each condition shown as a violin plot overlaid with 500 representative data points (n=4 cells per condition) (ns: not significant; ***: P<0.001, two-sample Kolmogorov-Smirnov test with Bonferroni correction for multiple comparisons). Not indicated in plot: Velocities of NiCl_2_-treated cells at 0 hours and 4 hours were significantly different (P<0.001).

As a functional test, we introduced a nickel chloride (NiCl_2_) solution into cage traps containing trapped cells. NiCl_2_ has been reported to inhibit the ciliary beating in *Stentor*, *Tetrahymena* and *Paramecium* by inhibiting axonemal dynein^6,14,35^. We measured cell motility for both untreated control cells and NiCl_2_-treated cells in an uncompressed trap at 0, 2, and 4 hours following media exchange. At each time point, NiCl_2_-treated cells were significantly slower than control cells (P<0.001, two-sample Kolmogorov-Smirnov test with Bonferroni correction for multiple comparisons) (Fig. 4c). The velocities of NiCl_2_-treated cells decreased significantly over time (P<0.001) (Fig. 4c). These results were expected as similar concentrations of NiCl_2_ have been reported to gradually inhibit ciliary beating in *Stentor* over the course of ∼5 hours^6^.

### Microsurgery of *Stentor* using SMORES

Finally, we demonstrate the utility of SMORES for microsurgical operations on *Stentor* by using different designs of the modular feature. To perform laser ablation of the cell membrane, we used the same modular feature (i.e., a confining ring) as that for immobilizing the cell. We immobilized the cell by compressing it to ∼30 μm in height and then ablated a user-defined region of interest. The laser ablation resulted in a cell membrane wound, indicated by a region with missing cortical pattern of pigment granules and km fibers (Fig. 5a). Without the immobilization provided by SMORES, the cell would swim away or contract rapidly to avoid the high-power laser used for ablation. The resulting wound, if inflicted at all, would be limited and would not correspond to the desired region of interest. Crucially, because SMORES released the compression slowly by lifting the cage trap away from the cell at a rate of ∼5 µm/second, we avoided further damage to the laser-ablated cell. Our design presents a major advantage over previous compression-based immobilization methods such as the microcompressor, which involves rotating a glass slide in contact with the cell^27,28^. We found that the microcompressor introduced high shear stress and further damaged the cell as we released the cell after ablation (Supplementary Video S4). In early attempts, we had investigated using a simple microchannel of gradually decreasing height to confine the cell. This approach had similar issues as the microcompressor, as flowing a laser-ablated cell while it was highly compressed resulted in high shear stress and further damaged the cell.

**Figure 5.**
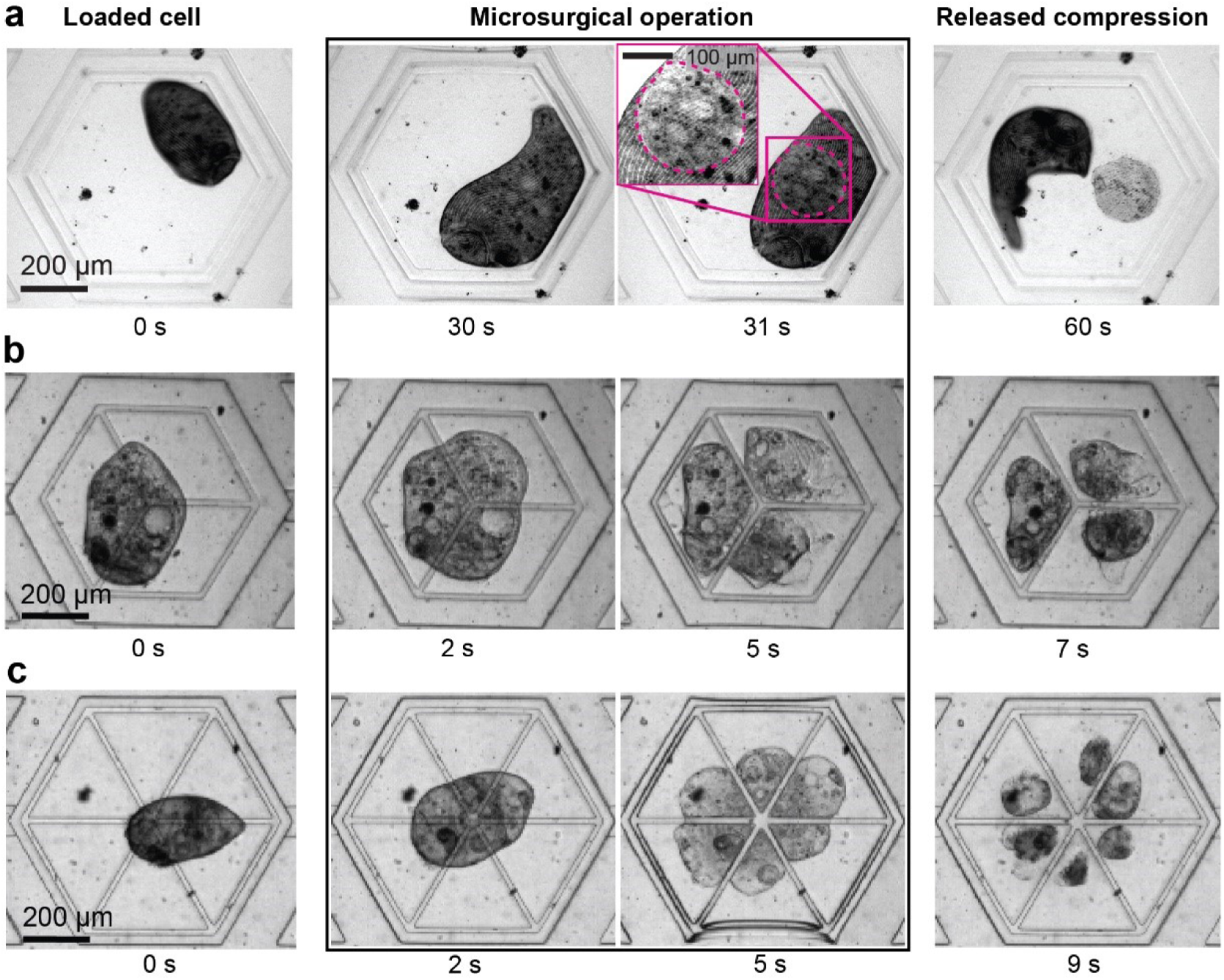
Microsurgery of *Stentor* using SMORES. (**a**) Laser ablation using the confining ring to immobilize the cell. Dashed line indicates membrane wound from laser ablation. Mechanical dissection of the cell into (**b**) 3 fragments using the 3-way cutter or (**c**) 6 fragments using the 6-way cutter.

In addition to laser ablation of the cell, we further demonstrated the utility of SMORES for the mechanical dissection of the cell by modifying the design of the modular feature to include a 3-way or 6-way cutter (Fig. 5b, c). Our objective was to demonstrate SMORES’ capability to perform complex cuts, a task challenging to achieve with conventional manual cutting using glass needles. We performed mechanical dissection by lowering the ceiling of the cage trap until the cutter contacted the glass substrate, effectively dissecting a trapped cell into three or six fragments simultaneously in ∼5 seconds (Fig. 5b, c). Similar to the SMORES operation for laser ablation, we could release the pressure in ∼2 – 5 seconds and retain the cells for observation within the cage trap. Alternatively, we could apply a flow in the flow layer to retrieve the dissected cells.

The working principle of our 3-way and 6-way cutters was similar to that of prior studies, in which a Quake valve lowered a PDMS pillar to create controlled wounds (i.e., gaps) in a cell monolayer^36^ or to perform axotomy of neurons^37^. However, considering the high motility of *Stentor*, the prior design would fail to wound *Stentor* as the cell would easily avoid a standalone PDMS pillar. Furthermore, our 3-way and 6-way cutters achieved more complex wounding patterns compared with the simple gaps used in prior designs.

## 3. CONCLUSIONS

Here we have presented a Simple Microfluidic Operating Room for the Examination and Surgery of *Stentor coeruleus* (SMORES). Compared with conventional Quake valve designs, our design achieves a more effective and uniform compression of a highly motile cell for up to 2 hours. The compression is reversible and minimally disruptive to the cell. The ease of media exchange allows long-term trapping and observation of a single cell, potentially over days or weeks. The ability to introduce drugs to the trapped cell and perform live cell imaging enables pharmacokinetic studies. Finally, the ability to perform laser ablation or mechanical dissection of a trapped cell allows one to monitor wound healing and regeneration processes not possible by hand surgery.

We designed our SMORES platform so that it can be used in laboratories without microfluidic infrastructure. Specifically, instead of using scientific syringe pumps, we opted for the use of a manually controlled 3D-printed pressure rig to control the compression on the cells. As the microfluidic channels of the SMORES platform occupies a total footprint of approximately 10 mm x 20 mm, the system has potential to be scaled to multiple traps on a single microfluidic chip in future work. Also, the device could be improved to further increase the uniformity of cell compression by further optimizing the design of the ceiling of the cage trap. Although we only tested our device on *Stentor coeruleus*, we believe that the design is directly applicable to other highly motile organisms, including other ciliates, and has the capacity to enable the investigation of biological processes previously not possible such as mechanotransduction and subcellular omics.

## 4. MATERIALS AND METHODS

### Device fabrication

#### Microfluidic devices

Microfluidic devices were made in PDMS (Sylgard-184, Dow) using standard soft lithography techniques^38,39^. We fabricated master molds in SU-8 resist: one mold each for the flow layer and the control layer. The flow layer required a multilayer mold containing resist features of multiple heights. We bonded the assembled PDMS devices on #1.5 coverslips with the flow layer seated directly against the coverslip to facilitate confocal imaging and laser ablation. This arrangement posed constraints for the PDMS replication process, as described below. The flow layer consisted of the cage trap whose ceiling acted as a deformable membrane (thickness ∼100 μm). During the fabrication of the PDMS replica on the mold for the flow layer, the presence of tall (up to ∼150 μm) resist features limited the uniformity or “flatness” of the top surface of the PDMS. This non-uniformity limited the use of conventional bonding by oxygen plasma to bond the flow layer to the control layer. Instead, we used a partial curing and bonding process^40^. The PDMS for all layers was formed using a base:crosslinker ratio of 10:1. The thickness of the PDMS on the mold for the flow layer was ∼100 μm above the tallest resist feature to form a PDMS membrane. The thickness of the PDMS on the mold for the control layer was ∼0.5 cm. After degassing, the PDMS on both molds was allowed to partially cure for ∼2 h at 70 °C. Next, the thick PDMS on the mold for the control layer was cut out, aligned, and placed on top of the thin PDMS on the mold for the flow layer. The two layers were then allowed to fully cure and crosslink together. Finally, the crosslinked control and flow layers were cut from the mold for the flow layer and then hole-punched and bonded to a #1.5 coverslip by oxygen plasma.

#### 3D-printed pressure rig

The 3D-printed pressure rig was designed in SolidWorks (Dassault Systèmes) and fabricated using Clear Resin on a Formlabs Form3 printer (Formlabs). An M3 x 40 bolt and M3 nuts were used as the leadscrew. The pressure rig was designed for a 3 mL syringe (Monoject).

##### Cell culture

*Stentor coeruleus* cells were cultured in Pasteurized Spring Water (PSW) (132458, Carolina Biological Supplies) in the dark at room temperature and fed *Chlamydomonas* algae every 2 days, as described previously^6^. All cells used for experiments were collected on the second day post-feeding.

##### Confocal microscopy

Confocal microscopy was performed on an inverted confocal scanning microscope (Zeiss, LSM 780) using Zen Black software (Zeiss).

##### Compression measurements

To visualize the compression of the cage trap, the cage trap was filled with 5 µM FITC in Pasteurized Spring Water (PSW) and imaged using a 488 nm laser and a 10X objective (NA 0.3). The pressure in the control layer was increased by advancing the leadscrew in the clockwise direction on the pressure rig until the cage trap just started to deform, and then the leadscrew was backed out slightly. This leadscrew position was marked as the initial leadscrew position. Defining the initial leadscrew position in this manner ensured the dampers were fully deformed before the start of the measurements, as the large dampers were designed to absorb initial pressure changes from the pressure rig. Consequently, further increase in pressure would contribute to deforming the cage trap only. The leadscrew was then incrementally advanced from the initial position to increase the cage trap deformation, and a z-stack of the cage trap was taken at each increment. Finally, the cage trap height was measured from each z-stack cross section using FIJI.

##### Live cell staining

A 1 mg/mL (3.14 mM) Nile Red stock solution was prepared by dissolving 1 mg of Nile Red (ThermoFisher, N1142) in 1 mL of DMSO. A 1 mM Vybrant DiD stock solution (ThermoFisher, V22887) and a 20 mM Hoechst 33342 stock solution (ThermoFisher, 62249) were purchased directly. Cells were first stained in 2 µg/mL (6.28 µM) Nile Red in PSW for 20 minutes, then washed in PSW. Cells were then stained in 2.5 µM DiD and 20 µM Hoechst 33342 in PSW for 20 minutes, then washed in 20 µM Hoechst 33342 in PSW. Finally, cells in 20 µM Hoechst 33342 in PSW were loaded into the cage traps and imaged immediately. We found that Hoechst did not stain well under the conditions tested, so we opted to leave it in the solution during imaging to maximize the fluorescence signal.

##### Laser ablation

Laser ablation was performed on an inverted confocal scanning microscope (Zeiss, LSM 780) using Zen Black software (Zeiss). Trapped cells were located and imaged using a 20X objective (NA 0.8), a 633 nm laser, and a transmitted light PMT (T-PMT) setup. After the cell was sufficiently compressed and immobilized, a region of interest was drawn, and ablation was performed using a 20X objective (NA 0.8) and a 561 nm continuous wave laser at 25% power (4.6 μW) and 25 μs exposure per pixel.

##### Data analysis

Cell centroids were extracted using OpenCV. Fluorescence intensity measurements were performed using FIJI. Statistical analyses were performed using SciPy, statsmodels, and NumPy in Python. Details of the statistical tests and sample size are provided in the corresponding text and figure legends.

## 6. ACKNOWLEDGEMENTS

We thank Prof. Wallace Marshall, Adrian Martin, Rebecca McGillivary, Ambika Nadkarni, and Rajorshi Paul for helpful discussions. This work was supported by the National Science Foundation (NSF Award: 1938109 and 2317442), and in part by the Center for Cellular Construction, which is a Science and Technology Center funded by the National Science Foundation (NSF Award: DBI-1548297). R.R. is supported by the NIGMS Center of the National Institutes of Health (Award: T32GM007276). Device fabrication was performed in the Stanford Nano Shared Facilities (SNSF), supported by the National Science Foundation (NSF Award: ECCS-2026822). Imaging was performed in the Stanford Cell Sciences Imaging Facility (CSIF) (RRID:SCR_017787).

## 7. AUTHOR CONTRIBUTIONS

K.S.Z. and S.K.Y.T. devised the concept for the work. K.S.Z., R.R., and S.K.Y.T. created device designs. K.S.Z. and R.R. fabricated devices. K.S.Z. and R.R. performed the experiments and data analysis. K.S.Z., R.R., and S.K.Y.T. wrote the manuscript. All authors read and approved the final manuscript.

## 9. DATA AVAILABILITY

The datasets generated and analyzed during the current study are available from the corresponding author on reasonable request.

## 5. REFERENCES

1. Ruehle, M. D., Orias, E. & Pearson, C. G. Tetrahymena as a unicellular model eukaryote: genetic and genomic tools. Genetics 203, 649–665 (2016).

2. Van Houten, J. Paramecium Biology. Results Probl Cell Differ 68, 291–318 (2019).

3. Howard-Till, R. A., Kar, U. P., Fabritius, A. S. & Winey, M. Recent advances in ciliate biology. Annu. Rev. Cell Dev. Biol. 38, 75–102 (2022).

4. Tartar, V. The Biology of Stentor. 5, (Pergammon, 1961).

5. Tang, S. K. Y. & Marshall, W. F. Self-repairing cells: How single cells heal membrane ruptures and restore lost structures. Science 356, 1022–1025 (2017).

6. Zhang, K. S., Blauch, L. R., Huang, W., Marshall, W. F. & Tang, S. K. Y. Microfluidic guillotine reveals multiple timescales and mechanical modes of wound response in *Stentor coeruleus*. BMC Biol. 19, 63 (2021).

7. Rajan, D. et al. Single-cell analysis of habituation in Stentor coeruleus. Curr. Biol. 33, 241–251.e4 (2023).

8. Tartar, V. Reactions of stentor coeruleus to homoplastic grafting. J. Exp. Zool. 127, 511–575 (1954).

9. Tartar, V. Grafting experiments concerning primordium formation in Stentor coeruleus. J. Exp. Zool. 131, 75–121 (1956).

10. Lin, A., Makushok, T., Diaz, U. & Marshall, W. F. Methods for the study of regeneration in stentor. J. Vis. Exp. (2018). doi:10.3791/57759

11. Lynn, D. H. The Ciliated Protozoa. (Springer Netherlands, 2008). doi:10.1007/978-1-4020-8239-9

12. Aufderheide, K. J. An overview of techniques for immobilizing and viewing living cells. Micron 39, 71–76 (2008).

13. Zhang, K. S., Nadkarni, A. V., Paul, R., Martin, A. M. & Tang, S. K. Y. Microfluidic surgery in single cells and multicellular systems. Chem. Rev. 122, 7097–7141 (2022).

14. Larsen, J. & Satir, P. Analysis of Ni(2+)-induced arrest of Paramecium axonemes. J. Cell Sci. 99 **( Pt** **1**), 33–40 (1991).

15. Tartar, V. Reactions of Stentor coeruleus to certain substances added to the medium. Exp. Cell Res. 13, 317–332 (1957).

16. Chokshi, T. V., Ben-Yakar, A. & Chronis, N. CO2 and compressive immobilization of C. elegans on-chip. Lab Chip 9, 151–157 (2009).

17. Rohde, C. B. & Yanik, M. F. Subcellular in vivo time-lapse imaging and optical manipulation of Caenorhabditis elegans in standard multiwell plates. Nat. Commun. 2, 271 (2011).

18. Chung, K., Crane, M. M. & Lu, H. Automated on-chip rapid microscopy, phenotyping and sorting of C. elegans. Nat. Methods 5, 637–643 (2008).

19. Marsot, P. & Couillard, P. The Use of Protamine-Coated Slides for Immobilizing Protozoa. J Protozool 20, 105–106 (1973).

20. Reize, I. B. & Melkonian, M. A new way to investigate living flagellated/ciliated cells in the light microscope: immobilization of cells in agarose. Botanica Acta 102, 145–151 (1989).

21. Jahn, T. L., Bovee, E. C. & Jahn, F. F. How to Know the Protozoa. 279 (W. C. Brown Company, 1979).

22. Soh, A. W. J. et al. Intracellular connections between basal bodies promote the coordinated behavior of motile cilia. Mol. Biol. Cell 33, br18 (2022).

23. Dong, L. et al. Reversible and long-term immobilization in a hydrogel-microbead matrix for high-resolution imaging of Caenorhabditis elegans and other small organisms. PLoS One 13, e0193989 (2018).

24. Le Berre, M., Aubertin, J. & Piel, M. Fine control of nuclear confinement identifies a threshold deformation leading to lamina rupture and induction of specific genes. Integr Biol (Camb) 4, 1406–1414 (2012).

25. Le Berre, M., Zlotek-Zlotkiewicz, E., Bonazzi, D., Lautenschlaeger, F. & Piel, M. Methods for two-dimensional cell confinement. Methods Cell Biol. 121, 213–229 (2014).

26. García-Arcos, J. M., Gateau, K., Venkova, L. & Piel, M. Extended methods for 2D confinement. Methods Mol. Biol. 2608, 63–81 (2023).

27. Yan, Y. et al. A microfluidic-enabled mechanical microcompressor for the immobilization of live single- and multi-cellular specimens. Microsc. Microanal. 20, 141–151 (2014).

28. Zinskie, J. A., Shribak, M., Bruist, M. F., Aufderheide, K. J. & Janetopoulos, C. A mechanical microcompressor for high resolution imaging of motile specimens. Exp. Cell Res. 337, 249–256 (2015).

29. Unger, M. A., Chou, H. P., Thorsen, T., Scherer, A. & Quake, S. R. Monolithic microfabricated valves and pumps by multilayer soft lithography. Science 288, 113–116 (2000).

30. Zeng, F., Rohde, C. B. & Yanik, M. F. Sub-cellular precision on-chip small-animal immobilization, multi-photon imaging and femtosecond-laser manipulation. Lab Chip 8, 653–656 (2008).

31. Guo, S. X. et al. Femtosecond laser nanoaxotomy lab-on-a-chip for in vivo nerve regeneration studies. Nat. Methods 5, 531–533 (2008).

32. Gokce, S. K. et al. A fully automated microfluidic femtosecond laser axotomy platform for nerve regeneration studies in C. elegans. PLoS One 9, e113917 (2014).

33. Okumus, B. et al. Mechanical slowing-down of cytoplasmic diffusion allows in vivo counting of proteins in individual cells. Nat. Commun. 7, 11641 (2016).

34. Okumus, B. et al. Single-cell microscopy of suspension cultures using a microfluidics-assisted cell screening platform. Nat. Protoc. 13, 170–194 (2018).

35. Bayless, B. A. et al. Asymmetrically localized proteins stabilize basal bodies against ciliary beating forces. J. Cell Biol. 215, 457–466 (2016).

36. Sticker, D., Lechner, S., Jungreuthmayer, C., Zanghellini, J. & Ertl, P. Microfluidic migration and wound healing assay based on mechanically induced injuries of defined and highly reproducible areas. Anal. Chem. 89, 2326–2333 (2017).

37. Hosmane, S. et al. Valve-based microfluidic compression platform: single axon injury and regrowth. Lab Chip 11, 3888–3895 (2011).

38. Xia, Y. & Whitesides, G. M. SOFT LITHOGRAPHY. Annu. Rev. Mater. Sci. 28, 153–184 (1998).

39. Duffy, D. C., McDonald, J. C., Schueller, O. J. & Whitesides, G. M. Rapid Prototyping of Microfluidic Systems in Poly(dimethylsiloxane). Anal. Chem. 70, 4974–4984 (1998).

40. Lai, A., Altemose, N., White, J. A. & Streets, A. M. On-ratio PDMS bonding for multilayer microfluidic device fabrication. J. Micromech. Microeng. 29, 107001 (2019).

