## Supplementary_information for "SMORES: A Simple Microfluidic Operating Room for the Examination and Surgery of *Stentor coeruleus*"

**Contents:**

**Supplementary Figure S1.** Additional details on the design of SMORES.

**Supplementary Figure S2.** Image processing pipeline.

**See online version for:**

**Supplementary Video S1.** Compressed *Stentor* cell inside a cage trap.

**Supplementary Video S2.** Uncompressed *Stentor* cell inside a cage trap.

**Supplementary Video S3.** Confocal z-stack of a *Stentor* cell.

**Supplementary Video S4.** Retrieval of a laser ablation-wounded cell from a microcompressor.

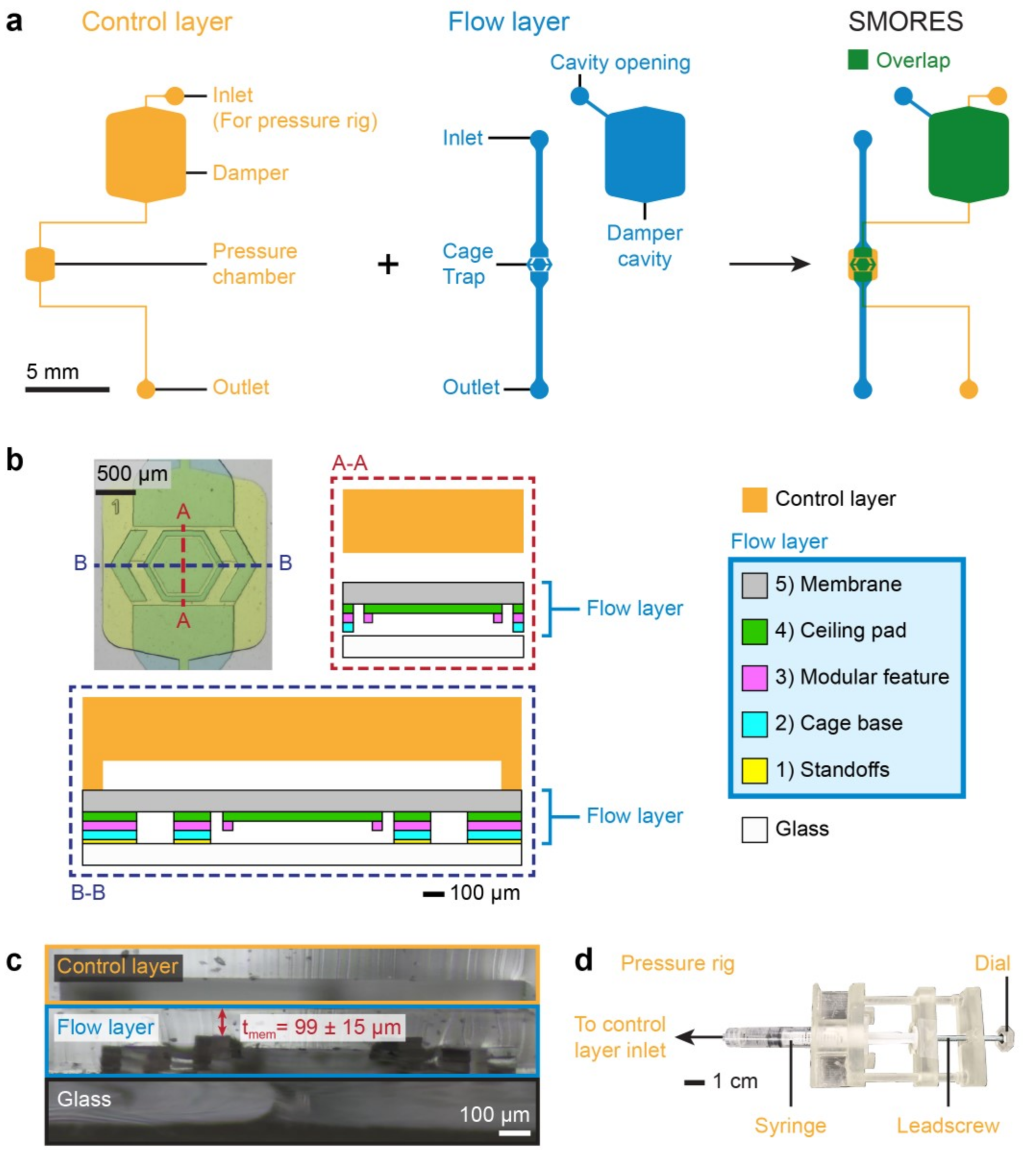


**Supplementary Figure S1.** Additional details on the design of SMORES. **(a)** Schematic diagram of the entire control layer and flow layer shown individually and assembled into the SMORES platform. **(b)** Layered components comprising the cage trap in the flow layer. Schematic diagrams of cross sections A-A and B-B correspond to the dashed lines in the top view shown in the top left. Layer heights are drawn to scale. **(c)** Cross-section view of a SMORES device that has been cut open approximately along the B-B section line. Membrane thickness $t_{mem}$ reported as mean ± standard deviation (n=15 devices). Note some distortion of the

PDMS features occurs due to the cutting. **(d)** 3D-printed pressure rig to interface with the control layer.

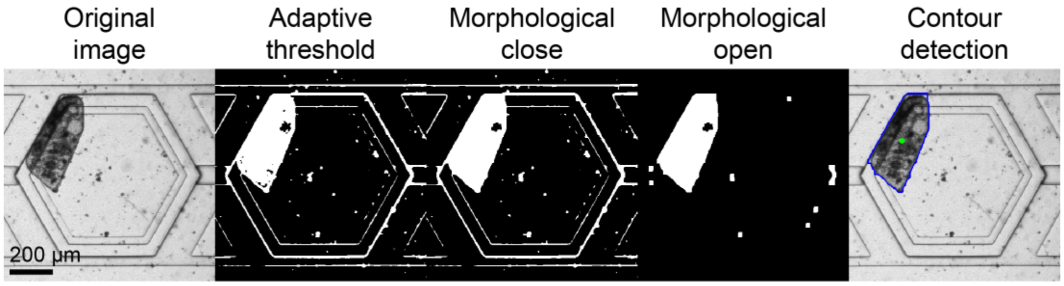


**Supplementary Figure S2.** Image processing pipeline. An adaptive threshold identifies dark regions in the original image. A morphological close reduces the noise present inside the cell body. A morphological open removes the thin borders of the cage trap features. Finally, contour detection locates the centroid of the cell. Note the image processing occasionally introduces artefacts into the detected cell contour; however, these artefacts do not change the results (also see Supplementary Videos S1 and S2).
